# The role of cation diffusion facilitator CDF-1 in lipid metabolism in *Caenorhabditis elegans*

**DOI:** 10.1101/2020.09.30.320986

**Authors:** Ying Hu, Yanli Wang, Xuanjun Wang, Xiaoyun Wu, Lin Fu, Xiayu Liu, Yu Wen, Jun Sheng, Jingjing Zhang

**Author notes:** These authors contributed equally to this work (Ying Hu, Yanli Wang). Corresponding authors, Corresponding authors: 1 Jingjing Zhang, Address: Chenggong Campus, Yunnan University South Section, East Outer Ring Road, Chenggong District, Kunming, Yunnan province, China, 2 Jun Sheng, Address: Yunnan Agricultural University, Heilongtan, North of Kunming, Kunming, Yunnan province, China.

## Abstract

Zinc is one of the most important trace elements that plays a vital role in many biological processes, and aberrant zinc metabolism has been implicated in lipid-related metabolic diseases. Previously, we showed that zinc antagonizes iron to regulate sterol regulatory element-binding proteins and stearoyl-CoA desaturase (SREBP-SCD) pathway in lipid metabolism in model organism *Caenorhabditis elegans*. Here, we further identified another cation diffusion facilitator CDF-1 in addition to SUR-7 in response to zinc to regulate lipid metabolism. Inactivation of SBP-1, the only homolog of SREBPs, leads to increased zinc level but decreased lipid accumulation reversely. However, either *cdf-1(n2527)* or *sur-7(tm6523)* mutation could successfully restore the altered fatty acid profile, fat content and zinc level of *sbp-1(ep79)* mutant. Furthermore, we found that CDF-1/SUR-7 may function bypass SBP-1 to directly affect the conversion activity of SCD in the biosynthesis of unsaturated fatty acids and lipid accumulation. Collectively, these results consistently support the link between zinc homeostasis and lipid metabolism via SREBP-SCD axis by cation diffusion facilitators CDF-1 and SUR-7.

## Introduction

Zinc is one of essential trace elements for living organisms, and plays numerous biological roles. For example, zinc determines the structural and functional of a variety of proteins, such as the metalloproteins, regulates both enzymatic activity and the stability of the proteins as either an activator or an inhibitor ion, and modulates cellular signal transduction processes that is usually regulated by zinc transport interrelated proteins, and so on (Fukada *et al*. 2011; Chasapis *et al*. 2012; Escobedo-Monge *et al*. 2019). Thus, it is critical and highly dynamic to maintain the zinc homeostasis for living organisms.

Dysregulation of zinc homeostasis is related with many human diseases. In particular, deficiency of zinc has been shown to be closely associated with a series of lipid metabolism diseases, including obesity and comorbid conditions such as insulin resistance, type 2 diabetes, inflammation, and altered lipid profile (Costarelli *et al*. 2010; Blazewicz *et al*. 2013; Miao *et al*. 2013). Erythrocyte zinc levels was associated with body mass index (BMI) and waist circumference(Blazewicz *et al*. 2013), and low zinc concentration was found in erythrocytes in obese women (Ennes Dourado Ferro *et al*. 2011). Moreover, there was a significant decrease in blood zinc levels in morbidly obese patients (de Luis *et al*. 2013). Meanwhile, zinc deficiency increases leptin production and exacerbates macrophage infiltration into adipose tissue in obese mice (Liu *et al*. 2013).

Zinc can be used as a nutrient to treat metabolic diseases. Zinc stimulates insulin secretion and increases sensitivity to insulin, and therefore zinc supplementation could improves insulin resistance, and improvement of blood pressure, glucose, and LDL cholesterol serum level in obesity (Cruz *et al*. 2017; Olechnowicz *et al*. 2018). Meanwhile, obesity-related cardiac hypertrophy (ORCH) could worsened by zinc deficiency (ZD) in HFD/ZD mice (de Luis *et al*. 2013), while reversely attenuated by zinc supplementary (ZS) in HFD/ZS group (Wang *et al*. 2016). However, it may have deleterious effects with excessive zinc supplementation, since excessive zinc intake may cause an undesirable increase in HbA1c levels and high blood pressure (Miao *et al*. 2013). Therefore, the balance of zinc in metabolism diseases is especially important.

Maintenance of intracellular zinc homeostasis depends on the balance of zinc absorption, distribution, and excretion, which is mainly regulated by two protein families metallothioneins (MTs) and zinc transport relevant proteins (Kimura and Kambe 2016). Zinc transport relevant proteins are mainly responsible for mediating the compartmentalization of zinc into various organelles and vesicles for their storage, and supply zinc to various proteins that require zinc for their function (Fukunaka and Fujitani 2018). Zinc transport relevant proteins are divided into two major families, the zinc transporters or cation diffusion facilitator (ZnTs/CDFs) or solute-linked carrier family 30 (SLC30A), and the Zip (Zrt- and Irt-like proteins) family or solute carrier 39A (SLC39A) (Liuzzi and Cousins 2004; Baltaci and Yuce 2018; Cotrim *et al*. 2019; Thokala *et al*. 2019). Several studies have shown that the gene expression of zinc transporters and disturbed zinc metabolism are associated with obesity and diabetes (Quraishi *et al*. 2005; Noh *et al*. 2014; Morais *et al*. 2019). The polymorphisms (SNP) of ZnT8 are in connection with type 1 (Xu *et al*. 2011; Wenzlau and Hutton 2013) and type 2 diabetes (Rutter and Chimienti 2015; Drake *et al*. 2017; Virgili *et al*. 2018). ZnT5 is also involved in metabolic diseases (Cuadrado *et al*. 2018). However, the underlying mechanisms of these zinc transporters in metabolic diseases are largely unknown.

Our previous work showed that SUR-7, one of zinc transport proteins in model organism *Caenorhabditis elegans* (Yoder *et al*. 2004; Roh *et al*. 2013), synergistically affects zinc homeostasis and lipid metabolism via SREBP-SCD axis (Zhang *et al*. 2017). In *C. elegans*, SBP-1 is the only homolog of sterol-regulatory element binding proteins (SREBPs), which are major regulators of lipid homeostasis including fatty acids, triglycerides, and cholesterol in vertebrate cells (Goldstein *et al*. 2006; Shao and Espenshade 2012). Inactivation of SUR-7 could restored the high level of zinc, low fat accumulation and fatty acid profiles of *sbp-1(ep79)* via directly upregulating the activity of stearoyl-CoA desaturase (Zhang *et al*. 2017). Thus, this raised a question whether other members of zinc transport related proteins, like SUR-7, were also involved in lipid metabolism. Thus, we examined zinc transport related proteins in *C. elegans*, and identified that *cdf-1(n2527)* mutation acted as another suppressor of *sbp-1(ep79)* mutant to restore its fat content and zinc levels, further providing consistent evidences to link zinc homeostasis and lipid metabolism.

## Materials and Methods

### Nematode strains, RNA interference (RNAi) and culture conditions

*C. elegans* strains were maintained on NGM plates with *E. coli OP50* under standard culture conditions, unless otherwise specified. RNAi was performed using the feeding method (Wu *et al*. 2018) and feeding bacterial strains was from the Ahringer *C. elegans* RNAi library. The wild type strain was Bristol N2 (WT). All the complete worm strains used in this study are listed in Table S1.

### Construct of *sur-7*RNAi clone

Construction of the *sur-7*RNAi were used *L4440* empty vector, the primers for the amplified empty vector and *sur-7* gene by PCR are shown in Table S2. These two linearized DNA fragments recombination followed the instruction of ClonExpress MultiS One Step Cloning Kit (Vazyme Biotech Co., Ltd), and then transformation into HT115 competent cells.

### Construction of transgenic strains

Construction of the *WT;kunEx203(cdf-1p::cdf-1::gfp)* and *WT;kunEx187(sur-7p::sur-7::gfp)* transgenic strains were done as followed. In general, DNA fragments of specific genes and related promoters were specific using PCR. The amplified DNA fragments were subsequently inserted into the transgenic plasmid *pPD95.75* (Frokjaer-Jensen *et al*. 2008). The injection mixture with 50 ng/μL transgenic plasmid and 5 ng/μ *L pCFJ90(Pmyo-2::mcherry)* and *rol-6(su1006)* were injected into the gonads of young adult wild type N2 worms. The positive transgenic worms were selected based on fluorescence expression. The primer sequences for the construction of transgenic worms were listed in Table S2.

### Nile Red staining of fixed worms and quantification of lipid droplet (LD) size

Nile Red staining of fixed worms was performed as previously described (Brooks *et al*. 2009; Liang *et al*. 2010). Young adult worms were collected and suspended in 1 mL of water on ice, then resuspended in 1 mL of buffer M9 and 50 μL of freshly prepared 10% paraformaldehyde solution. Worms were frozen in liquid nitrogen, subjected to two or three incomplete freeze/thaw cycles, and then washed with M9 buffer several times to remove the paraformaldehyde solution. Two microliters of 5 mg/mL Nile Red was added to the worm pellet (1 mL) and incubated for 30 min at 20°C, with occasional gentle agitation. Worms were washed two or three times with M9 buffer and mounted onto 2% agarose pads for microscopic observation and photography. For each animal, a projection image with proper intensity was acquired using an Olympus BX53 fluorescence microscope (Japan) (Shi *et al*. 2013). At least 15 animals were imaged and lipid droplet size quantified approximately 5 worms for each worm strain or treatment.

### Analysis of fatty acid compositions

Approximately 2,000 young adults were harvested for fatty acid exterification and analysis. The harvested worms’ fatty acids were esterified with 1 mL of MeOH + 2.5% H_2_SO_4_, heated 1 hour under 70°C. Determination of the fatty acids were run with an Agilent 7890 series gas chromatographer equipped with a 15 m × 0.25 mm DB-WAX column (Agilent, USA), with helium as the carrier gas at 1.4 mL/min, and a flame ionization detector. Four biological replicates were performed for GC analysis.

### Analysis of triacylglycerol (TAG)

About 4×10^4^ synchronized young adults were harvested for total lipid extraction, the separation of triacylglycerol (TAG) and phospholipids (PL) by thin-layer chromatography (TLC) and the determination of fatty acids by gas chromatography (GC). Fatty acid compositions were analyzed with an Agilent 7890A, and methyl ester of C15:0 was used as a standard for quantitative analysis.

### Supplementation or sequestration of zinc

ZnSO_4_ supplementation and zinc reduction by N,N,N’,N’-tetrakis (2-pyridylmethyl) ethylenediamine (TPEN) were performed as described previously (Zhang *et al*. 2017). In brief, synchronized L1 worms were placed and cultivated on NGM plates supplied with either ZnSO_4_ or TPEN with a final concentration of 50 and 5 μM respectively, and young adult worms were harvested for further analysis.

### Zinpyr-1 staining and visualization

Zinpyr-1 staining was performed as described previously (Roh *et al*. 2013). The fluorescence of Zinpyr-1 was visualized under an OLYMPUS BX53 fluorescence microscope (Olympus, Japan). Images were captured using identical settings and exposure times, unless specifically noted. The fluorescence intensity was quantified using Photoshop software (Blazewicz *et al*. 2013).

### Visualization of GFP

L4 and young adult GFP positive worms were picked and plated on an agarose gel pad and visualized under Two-photon confocal microscope (Nikon A1MP+) or a fluorescent microscope (BX53, Olympus, Japan). Images were captured using the same settings and exposure times for each worm, unless specifically indicated, and the GFP reporter expression was quantified using Photoshop software.

### Growth Rate assay

The gravid adult worms were treated with alkaline hypochlorite and obtained their eggs, which were then seeded onto NGM plates, cultured for several days until they grown up to adults. The number of adults and the total number of worms were scored under a microscope at each 12 hours after seeded for 48 hours, each strain underwent three biological replications.

### Data analysis

Data are presented as the mean ± SEM, except when specifically indicated. Statistical analysis was conducted using Student’s t-test. Figures were made using GraphPad Prism 6 (GraphPad Software, La Jolla, CA, USA) and Adobe Illustrator CS6.

## Results

### The cation diffusion facilitator CDF-1 maintained zinc homeostasis and lipid metabolism

In eukaryotic cells, two major families of zinc transport proteins, cation diffusion facilitators (CDF/ZnT/SLC30) and zrt-, irt-like proteins (ZIP/SLC39)), mediate zinc trafficking and homeostasis. In total, 14 CDFs and 14 ZIPs in *C. elegans* have been identified with the homologous protein sequences to the 10 CDF and 14 ZIP proteins in *Homo sapiens* (Kambe *et al*. 2015) (Figure S1). We previous reported one of above zinc transport proteins SUR-7 that coordinately affects zinc homeostasis and lipid metabolism (Zhang *et al*. 2017). To explore whether other related transporters also involved in zinc homeostasis and lipid metabolism, we initially tested 11 out of 28 zinc related transporters with available mutants to detected their labile zinc and pseudocoelomic zinc by zinpyr-1 fluorescence and lipid droplets by Nile Red staining, respectively (Brooks *et al*. 2009; Roh *et al*. 2013).

Zinpyr-1 staining showed differential intensity and location of fluorescence among various worm mutants under the treatment with or without ZnSO_4_. (Figure 1A and B, Figure S2). Under normal condition, like *sur-7(tm6523)* mutant, *cdf-1(n2527), zipt-2.3(ok2094)*, and *zipt-15(ok2160)* mutants displayed reduced fluorescence (Figure 1A and B, Figure S2), which was mainly restricted in the area of intestine lumen, suggesting an defect in uptake of zinc. In contrast, the majority of mutants especially the *ttm-1(tm6669)* mutant displayed higher zinpyr-1 fluorescence than the wild type N2 (Figure 1A and B, Figure S2). In the presence of ZnSO_4_, most of worm mutants of zinc related transporters showed an increased level of Zinpyr-1 fluorescence to some extent (Figure 1A and B, Figure S2). Interestingly, among 11 mutants of zinc related transporters, only *cdf-1(n2527)* mutant displayed reduced lipid droplet size and lipid accumulation similar to *sur-7(tm6523)* mutant in response to the treatment of ZnSO_4_. However, the lipid droplet (LD) size and lipid accumulation of the other worms were not altered under normal condition (Figure 1C and D, Figure S3). Furthermore, consistent to our previous report (Zhang *et al*. 2017), reduction of zinc by TPEN (N,N,N’,N’-Tetrakis (2-pyridylmethyl) ethylenediamine), a zinc chelator, led to significantly increased LD size and lipid accumulation in all tested mutants of zinc related transporters like N2 (Figure 1C and D, Figure S3). Taken together, these results indicated that CDF-1 may function like SUR-7 in regulating zinc homeostasis and lipid metabolism.

**Figure 1.**
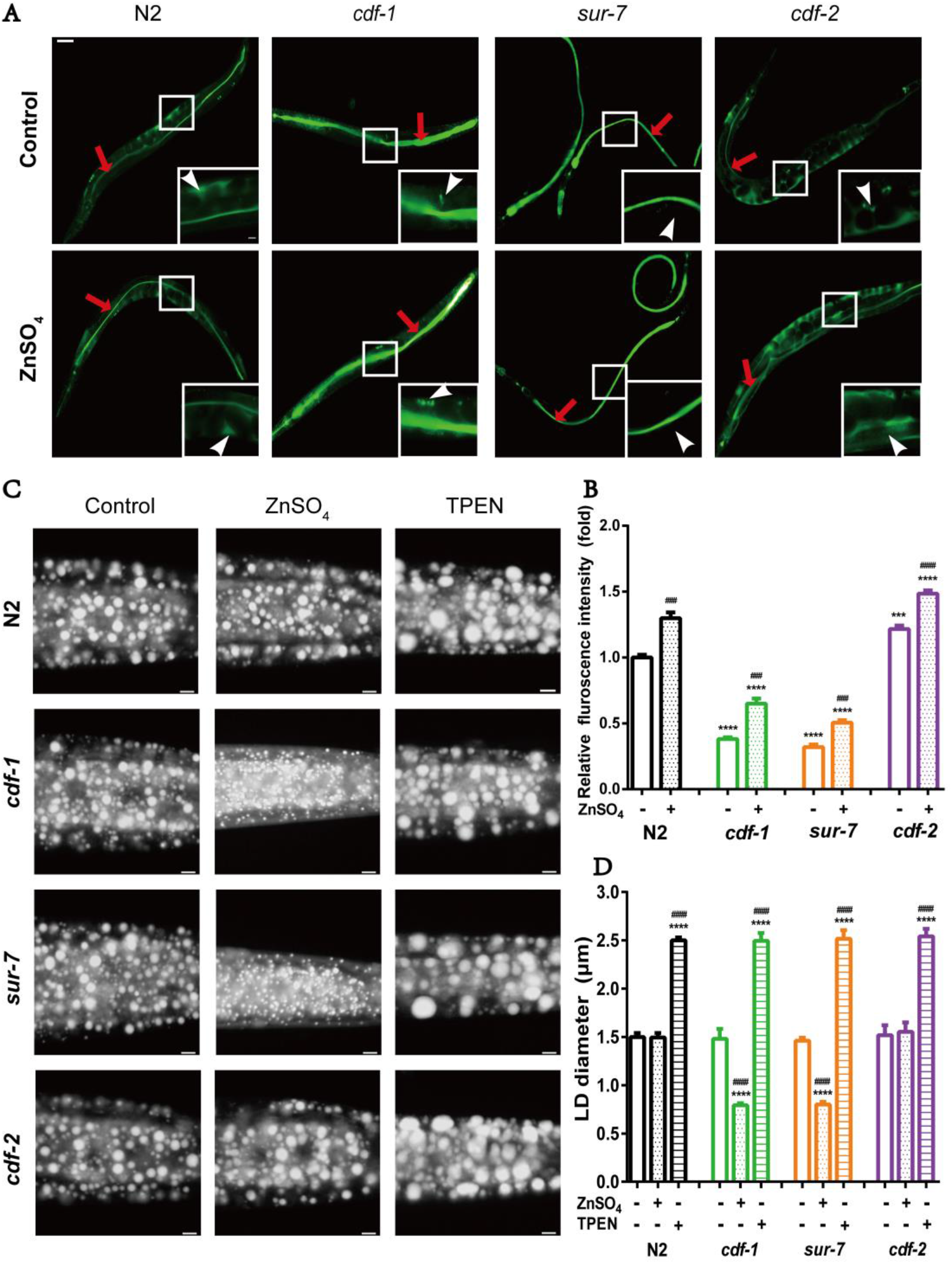
The mutants of zinc transporters with zinc level and fat accumulation. (A) Fluorescence microscopy of WT, *cdf-1(n2527), sur-7(tm6523)* and *cdf-2(tm788)* treated with or without ZnSO_4_, which stained with Zinpyr-1 and captured using identical settings and exposure times. Scale bar represents 50 μm for whole worms and 10 μm for enlarged worms. The concentration of ZnSO_4_ is 50 μM. The red arrowheads indicate the lumen zinc, the white arrows show pseudocoelomic zinc. (B) Quantitation of the fluorescence intensity about pseudocoelomic zinc from (A). (C) Nile red staining of fixed worms treated with ZnSO_4_ or TPEN. For representative animals, the anterior was on the left and the posterior was on the right, Bar, 5 μm. (D) Quantitation of lipid droplet size in the posterior region of intestines from five worms of each worm strain. The data are presented as mean ± SEM. For all statistics analysis, significant difference between WT N2 and a specific condition: ***P < 0.001, ****P < 0.0001(unpaired t-test). Significant difference between the control and ZnSO_4_/TPEN treatment of each strain: ^###^P < 0.001, ^####^P < 0.0001 (unpaired t-test).

### Inactivation of CDF-1 restored the lipid metabolism of *sbp-l(ep79)* mutant

Our previous study showed that SUR-7 functions *via* the sterol-regulatory element binding protein (SREBP) and its target stearoyl-CoA desaturase (SCD) to regulate lipid metabolism (Zhang *et al*. 2017). Inactivation of *sur-7* restores the lipid metabolism of *sbp-1(ep79)* mutant by upregulating the activity of SCD. As mentioned above, the *cdf-1(n2527)* mutant displayed similar phenotypes like *sur-7(tm6523)* mutation in responses to zinc, therefore, we wondered whether CDF-1 also affected lipid metabolism *via* the SREBP-SCD axis.

SUR-7, CDF-1 and CDF-2 proteins belong to the CDF family, and they possess six conservative transmembrane domains and (Figure S1 and S4). The *cdf-1(n2527)* mutation contains a SNP in the third transmembrane domain, leading to a premature termination (Figure S4A). The *cdf-2(tm788)* mutation contains a 804 bp deletion and 68 bp insertion, which cause frame shift mutation (Figure S4B). The *sur-7(tm6523)* mutation contains a 564 bp deletion and 4 bp insertion, which cause early termination of translation (Figure S4C). To verify whether *cdf-1* could be a suppressor of *sbp-1(ep79)*, we constructed the *sbp-1(ep79);cdf-1(n252 7)* double mutants, and *sbp-1(ep79);sur-7(tm6523)* was used as positive control, while the *sbp-1(ep79);cdf-2(tm788)* was used as negative control. As we expected, the LD size and triacylglycerol (TAG) content of both *sbp-1(ep79);cdf-1(n2527)* and *sbp-1(ep79);sur-7(tm6523)*, but not the *sbp-1(ep79);cdf-2(tm 788)* mutants were significantly increased compared to the *sbp-1(ep79)* mutant (Figure 2A-C). Similarly, the *sbp-1(ep79);cdf-1(n2527)* worms developed to adulthood about 24 hours earlier than the *sbp-1(ep79)* worms (Figure 2D). Taken together, these results demonstrated that the *cdf-1(n2527)* mutation was another suppressor of *sbp-1(ep79)* mutant, in which it could restore the lipid accumulation and growth rate of *sbp-1(ep79)* mutant.

**Figure 2.**
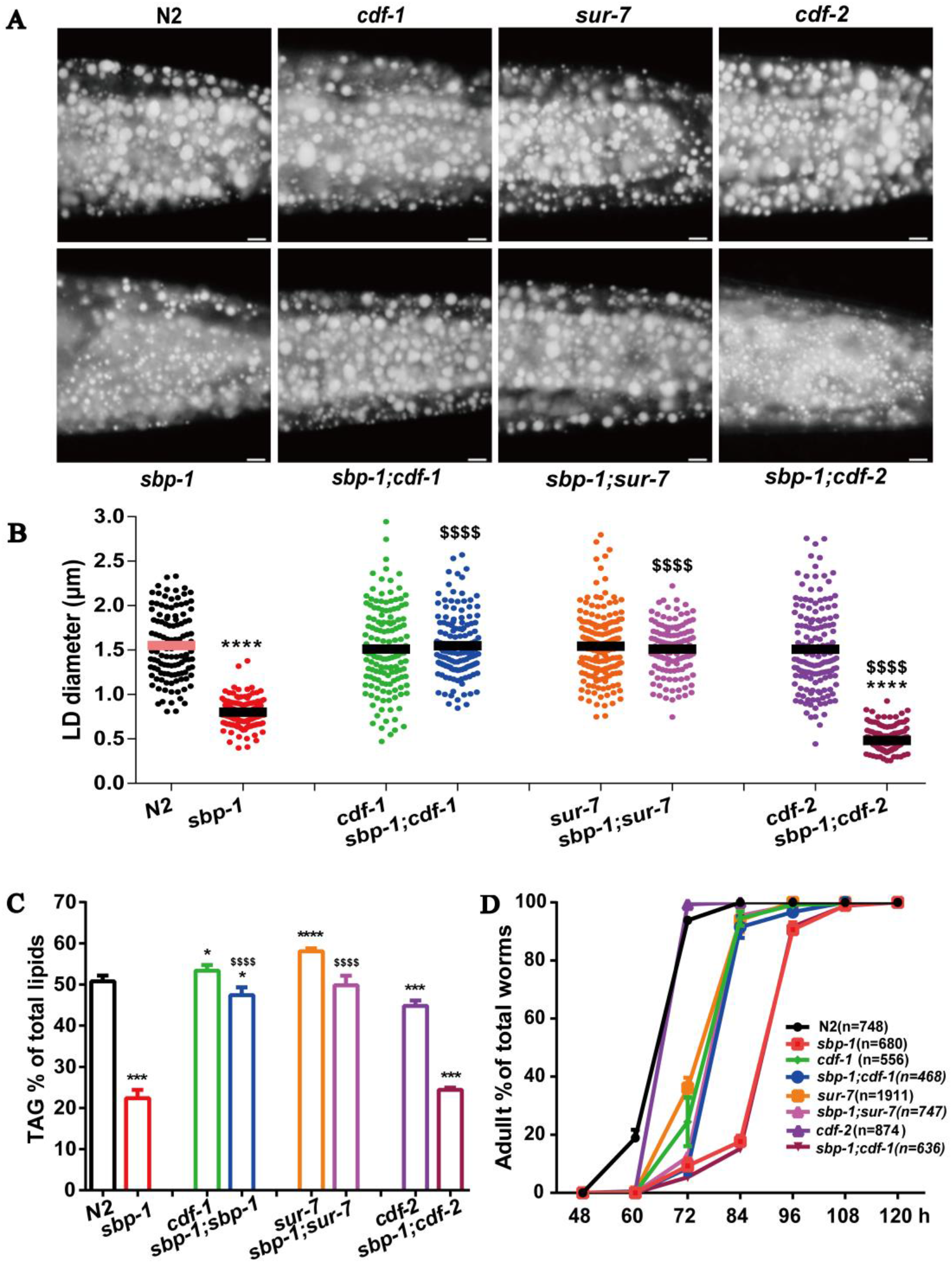
*cdf-1* and *sur-7* restored lipid profiles of *sbp-1*. (A) The lipid droplets by Nile red staining in the worms of WT, *sbp-1(ep79)*, *cdf-1(n2527)*, *sbp-1(ep79;cdf-1(n2527)), sur-7(tm6523), sbp-1(ep79);sur-7(tm6523), cdf-2(tm788)* and *sbp-1(ep79);cdf-2(tm788)*. For representative animals, the anterior was on the left and the posterior was on the right, Bar, 5 μm. (B) Quantitation of lipid droplet diameters in the posterior region of intestines from five worms of each worm strain. The data are presented as mean ± SEM. (C) The percentage content of triacylglycerol (TAG %) in total lipids (TAG+PL, phospholipids) of each worm strain. The data are presented as mean ± SEM of four biological repeats. (D) The growth rate of worms. The data are presented as the mean ± SEM with three biological repeats, n: the number of scored worms for each strain. For all statistics analysis, significant difference between WT N2 and each mutant: *P < 0.05, **P < 0.01, ***P < 0.001, ****P < 0.0001. Significant difference between *sbp-1(ep79)* and double mutants: ^$^P < 0.05, ^$$^P < 0.01, ^$$$^P < 0.001, ^$$$$^P < 0.0001.

### The suppression of *sbp-1(ep79)* by *cdf-1(n2527)* was related to zinc level

As our previous study demonstrated, the *sbp-1(ep79)* mutant displays increased level of Zinpyr-1 fluorescence but reduced lipid accumulation, which can be reversed by *sur-7(tm6523)* (Zhang *et al*. 2017). Consistently, like the *sbp-1(ep79);sur-7(tm6523)* mutant, the zinc level indicated by Zinpyr-1 fluorescence of the *sbp-1(ep79);cdf-1(n2527)* mutants was significantly reduced compared with the *sbp-1(ep79)* worms, even under the treatment of ZnSO_4_ (Figure 3A and B), suggesting the upregulated zinc level of *sbp-1(ep79)* mutant depends on the activity of SUR-7 and CDF-1.

**Figure 3.**
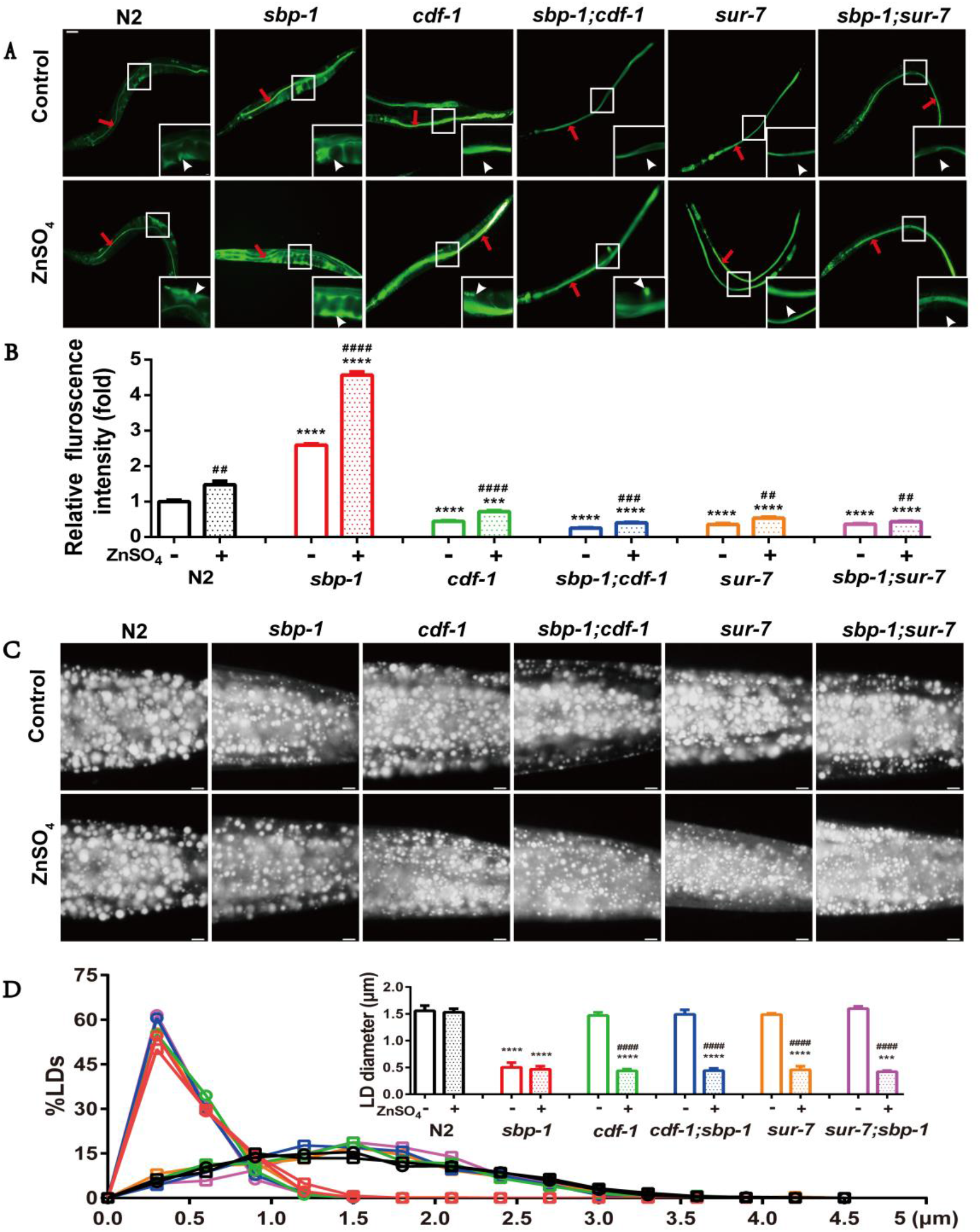
Zinc regulated fat accumulation in *C. elegans*. (A) zinc levels of WT, *sbp-1(ep79), cdf-1(n2527), sbp-1(ep79;cdf-1(n2527)), sur-7(tm6523)*, *sbp-1(ep79);sur-7(tm6523), cdf-2(tm788)* and *sbp-1(ep79);cdf-2(tm 788)* treated with or without ZnSO_4_ by stained with Zinpyr-1. Images were captured using identical settings and exposure times. Scale bar represents 50 μm for whole worms and 5 μm for enlarged worms. The concentration of ZnSO_4_ is 50 μM. The red arrowheads indicate the lumen zinc accumulates of the intestine, the white arrows show pseudocoelomic zinc. (B) Quantitation of the fluorescence intensity. (C) Nile red staining of fixed worms’ lipid droplets, Bar, 5μm. (D) Quantitation of lipid droplet size and frequency distribution of LD diameter in the posterior region of intestines from five worms of each worm strain. All of quantitation of the data is presented as mean ± SEM. For all statistics analysis, significant difference between N2 (WT) and a specific condition: *P < 0.05, **P < 0.01, ***P < 0.001, ****P < 0.0001. Significant difference between within groups of the control and ZnSO_4_ treatment: ^#^P < 0.05, P < 0.01, ^###^P < 0.001, ^####^P < 0.0001 (unpaired t-test), which applies to all subsequent statistical analysis.

As we mentioned above and previous work reported (Zhang *et al*. 2017), zinc negatively regulates lipid accumulation, and zinc reduction leads to increased lipid accumulation. Next, we asked whether the restoration of lipid accumulation of *sbp-1(ep79)* mutant by *cdf-1(n2527)* was related to zinc level. Nile Red staining of fixed worm showed that the LD sizes of *sbp-1(ep79);cdf-1(n2527)* and *sbp-1(ep79);sur-7(tm6523)* mutants were significantly reduced to the level of *sbp-1(ep79)* mutant when treated with ZnSO_4_, similar to the *cdf-1(n2527)* or *sur-7(tm6523)* single mutant alone (Figure 3C and D). However, the LD size in N2 and *sbp-1(ep79)* mutant were not changed between ZnSO_4_ treatment and not (Figure 3C and D). Altogether, these results further support a tight connection of lipid accumulation and zinc homeostasis regulating by SREBP-CDF-1/SUR-7.

### CDF-1 regulated the activity of SCD required zinc

In *C. elegans*, the FAT-5, FAT-6 and FAT-7 are three SCDs that convert saturated fatty acids palmitic acid C16:0 and stearic acid C18:0 to palmitoleic acid C16:1(n-7) and oleic acid C18:1(n-9) (Brock *et al*. 2007; Shi *et al*. 2013; He *et al*. 2018). Our previous study showed that the dietary ZnSO_4_ reduction of lipid accumulation in *sur-7(tm6523)* mutant was due to a decreased conversion activity of SCD (Zhang *et al*. 2017). To conform the SCD conversion activity in the mutants contented *cdf-1(n2527)*, we detected their fatty acid profiles *via* gas chromatography (GC) under either TPEN or ZnSO_4_ treatment. The levels of oleic acid C18:1(n-9) and the conversion activity of SCD (C18:1(n-9)/C18:0) were increased when treated with TPEN in all worm strains (Figure 4A-C), while they were reduced in *cdf-1(n2527)* and *sur-7(tm6523)* mutants in the presence of ZnSO_4_ (Figure 4A-C). Meanwhile, the levels of saturated fatty acids (C16:0 and C18:0) in *sbp-1(ep79);cdf-1(n2527)* and *sbp-1(ep79);sur-7(tm6523)* mutants were reduced compared with *sbp-1(ep79)* mutant (Figure 4A and D). Consistently, the SCD conversion activity was increased of *sbp-1(ep79);cdf-1(n2527)* and *sbp-1(ep79);sur-7(tm6523)* mutants (Figure 4C and F). Collectively, these results indicated that the *cdf-1(n2527)* mutant was similar to *sur-7(tm6523)* mutant in response to dietary ZnSO_4_ or TPEN during regulating SCD activity.

**Figure 4.**
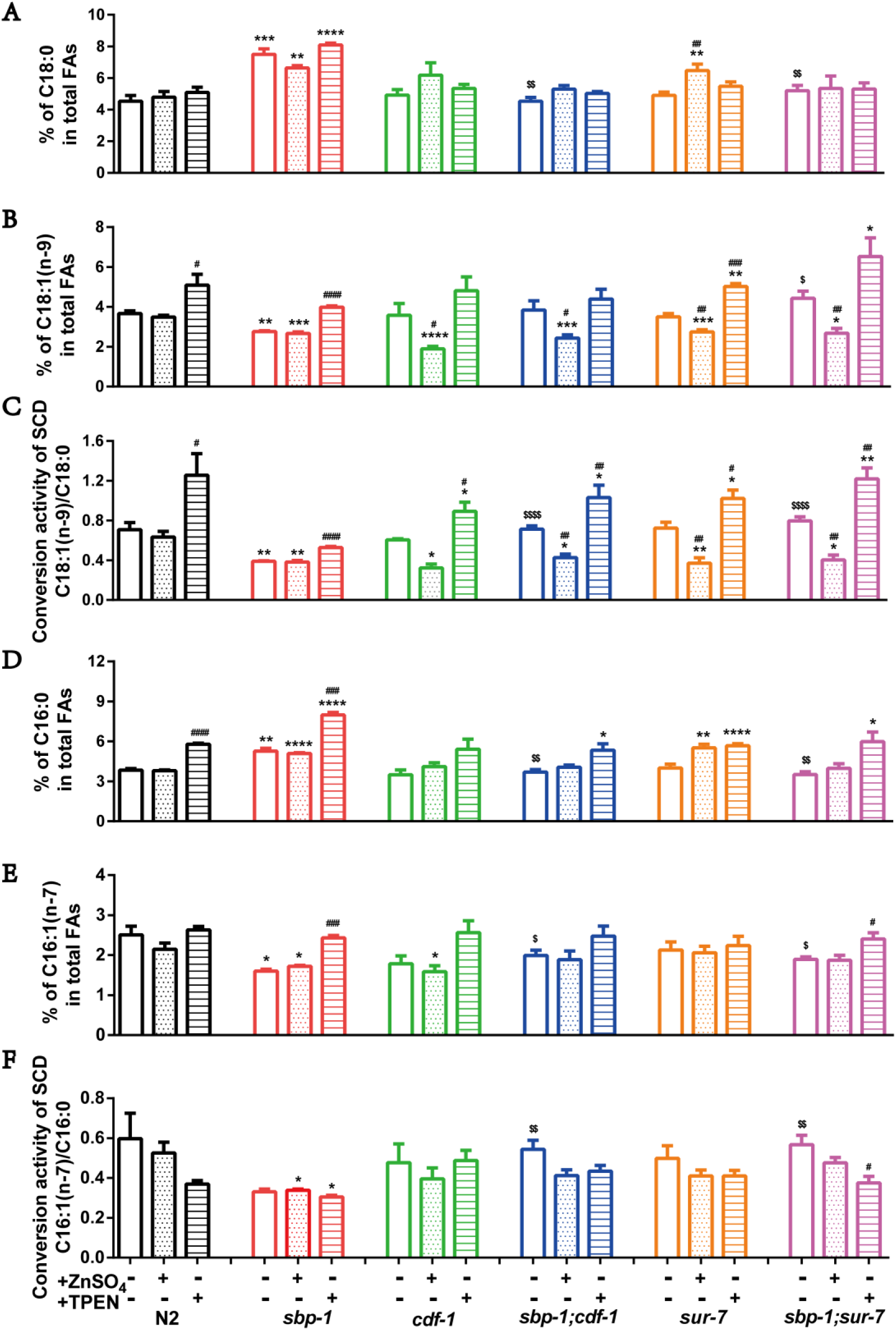
*cdf-1* and *sur-7* mediated the activity of SCD by zinc. (A), (B), (D) and (E) The fatty acid profiles of N2 (WT), *sbp-1(ep79)*, *cdf-1(n2527)*, *sbp-1(ep79;cdf-1(n2527))*, *sur-7(tm6523)* and *sbp-1(ep79);sur-7(tm6523)* were quantified using GC analysis. (A) and (B) Percentage of C18:0 and C18:1(n-9) in total fatty acids. (C) The conversion activity of SCD presented by the ratio of C18:1(n-9)/C18:0. (D) and (E) Percentage of C16:0 and C16:1(n-7) in total fatty acids. (F) The conversion activity of SCD presented by the ratio of C16:1(n-7)/C16:0. The data are presented as mean ± SEM of more than three biologic repeats.

### Zinc promotes SCD expression but reversely inhibits its conversion activity

Since the *n2527* mutation of *cdf-1* affects the SCD activity under ZnSO_4_ treatment, we therefore ascertain whether it was due to the altered expression of SCDs. In order to test the expression of SCDs, we opted to use *cdf-1*RNAi and *sur-7*RNAi for experimental convenience. Meanwhile, we constructed two transgenic strain, *WT;kunEx203(Pcdf-1::cdf-1::gfp)* and *WT;kunEx187(Psur-7::sur-7::gfp)*. The CDF-1::GFP is mainly expressed in intestine and SUR-7::GFP is mainly expressed in muscle. The *cdf-1*RNAi specifically inhibited the fluorescence expression of CDF-1::GFP but not SUR-7::GFP. Likewise, The *sur-7*RNAi only inhibited the fluorescence expression of SUR-7::GFP but not CDF-1::GFP, suggesting the efficiency and specification of each gene RNAi (Figure S5). The fluorescence intensity of FAT-5::GFP, FAT-6::GFP and FAT-7::GFP were not changed in *cdf-1*RNAi and *sur-7*RNAi worms compared to the control (empty vector, EV) (Figure 5). Interestingly, distinguished from the SCD conversion activity (Figure 4C), the fluorescence intensity of FAT-5::GFP, FAT-6::GFP and FAT-7::GFP was significantly increased in the presence of ZnSO_4_, but was decreased under zinc reduction by TPEN treatment in control (EV), *cdf-1* RNAi and *sur-7*RNAi worms (Figure 5), which were contradictory with the SCD conversion activity reduced under ZnSO_4_ or increased TPEN treatment. Nevertheless, these results suggest that zinc promotes SCD expression but reversely inhibits its conversion activity.

**Figure 5.**
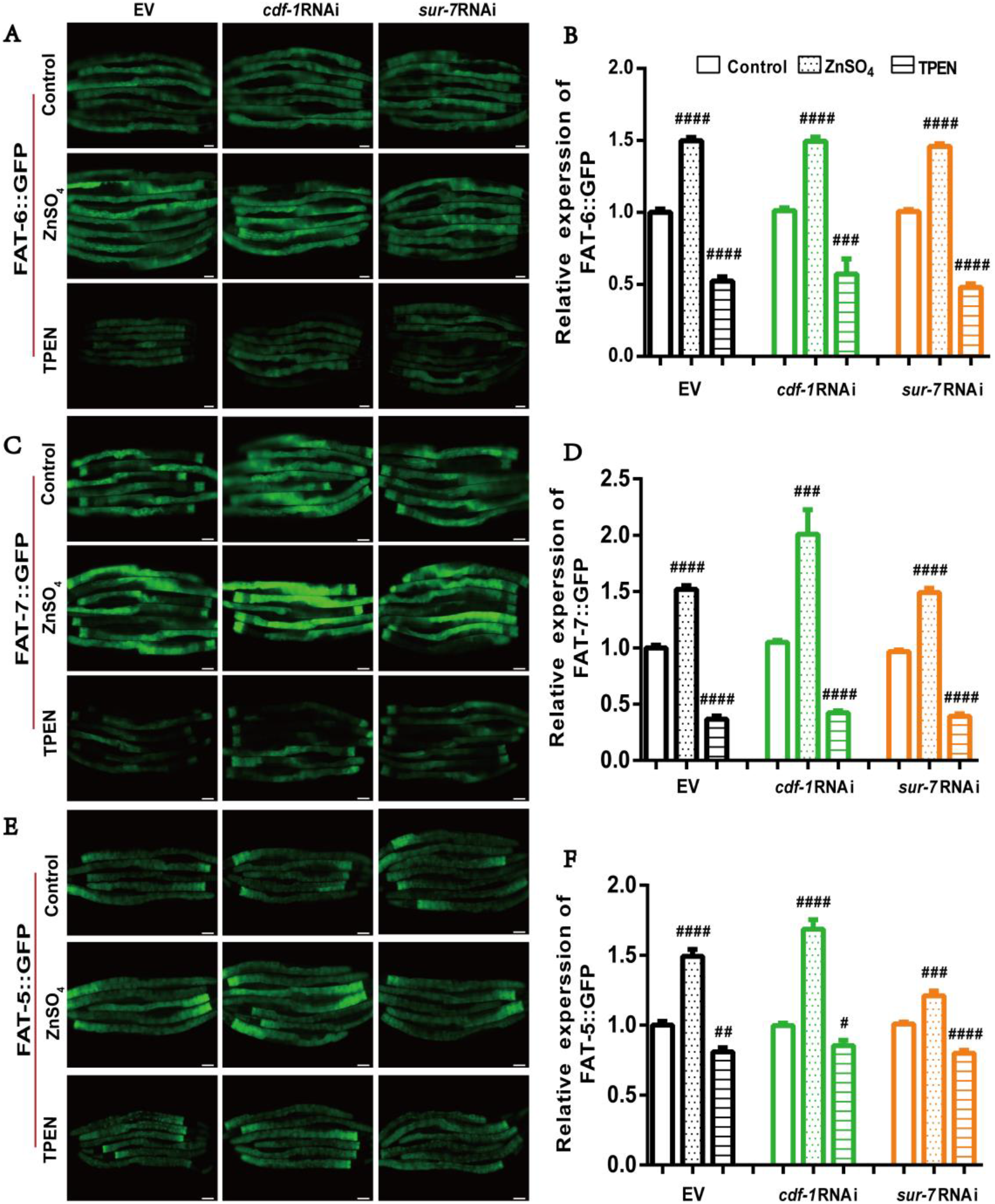
The expression of FAT-6::GFP, FAT-7::GFP and FAT-5::GFP under empty vector or a specific condition. (A), (C) and (E) The fluorescence intensity of FAT-6::GFP, FAT-7::GFP and FAT-5::GFP. Bar, 50 μm. From left to right were EV, *cdf-1*RNAi and *sur-7*RNAi, from top to bottom were control, ZnSO_4_ supplementation and TPEN supplementation. (B), (D) and (F) The quantization of the GFP fluorescence intensity corresponding to figure (A), (C), (E), respectively.

## Discussion

Zinc is an essential element for living organisms. Dysfunction of zinc homeostasis is associated with lipid metabolism-related diseases. Our previous work uncovered a cation diffusion facilitator SUR-7 that is required for the function of SREBP-SCD in lipid synthesis and accumulation in *C. elegans*. In this study, we further identified another cation diffusion facilitator CDF-1 among other zinc related transporters that functions similar to SUR-7 in regulating lipid metabolism. We proposed a model to better illustrate the function of CDF-1/SUR-7 in lipid metabolism. Under normal condition, CDF-1/SUR-7 may involve in zinc uptake and transportation for maintenance of zinc homeostasis as well as down regulation of fatty acid biosynthesis and lipid accumulation. Inactivation of CDF-1 or SUR-7 impairs zinc uptake and transportation, which result in reduction of zinc that consequently upregulate the conversion activity of stearoyl-CoA desaturase (SCD), further promoting the biosynthesis of unsaturated fatty acids and lipid accumulation (Figure 6).

**Figure 6.**
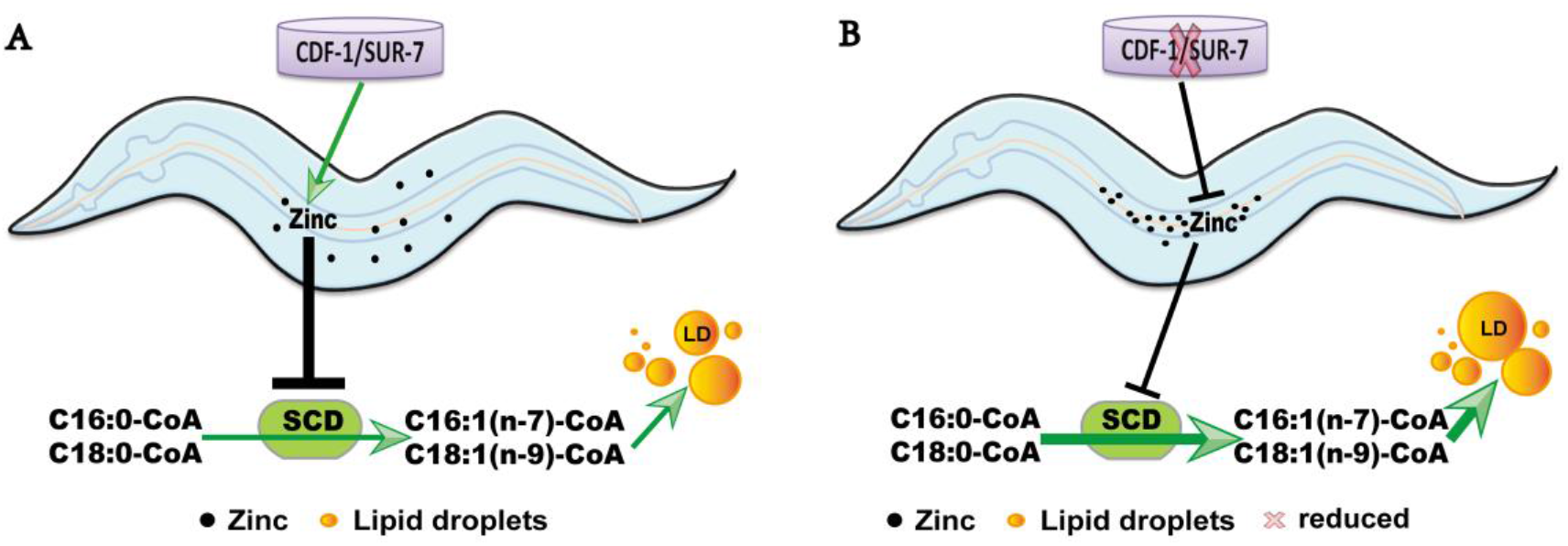
A working model of CDF-1 and SUR-7 mediate zinc to regulate SCD activity in lipid metabolism. (A) Under normal condition, CDF-1 and SUR-7 promote the transport of zinc into the worm body, which prevented SCD from over activation to maintain the biosynthesis of MUFAs and fat accumulation. (B) As CDF-1 and SUR-7 were impaired, zinc transporting into the worm body from lumen was blocked, which enhanced the activity of SCD and promoting the biosynthesis of MUFAs and accelerating fat accumulation.

Zinc transport relevant proteins play crucial roles in maintaining of zinc homeostasis. Dysfunction of zinc transporters is associated with obesity and diabetes (Quraishi *et al*. 2005; Noh *et al*. 2014; Morais *et al*. 2019). In *C. elegans*, a total of 28 zinc transport relevant proteins have been identified based on the protein sequences to human homologues (Kambe *et al*. 2015). Among these zinc transport proteins, we found that CDF-1 and SUR-7 function redundantly to regulate zinc homeostasis and lipid metabolism. Inactivation of either CDF-1 or SUR-7 reduced Zinpyr-1 fluorescence in wild type N2 and also *sbp-1(ep79)* mutant. More importantly, both gene mutations could recover the LD size of *sbp-1(ep79)* mutant, and displayed similar response with reduction of LD size to ZnSO_4_ treatment. Therefore, our current work in *C. elegans* also supports a negative relationship between zinc level and lipid accumulation.

Stearoyl-CoA desaturase (SCD) is a main target of SREBP. SCD converts saturated fatty acids (C16:0 and C18:0) to MUFAs [C16:1(n-7) and C18:1(n-9)] for the biosynthesis of TAGs, phospholipids, and cholesterol esters. The SCD catalysis is dependent on its di-iron center. We previously showed that zinc reduction led to iron overload, consequently activating the conversion activity of SCD and fat accumulation (Zhang *et al*. 2017). Although the expression of FAT-5::GFP, FAT-6::GFP and FAT-7::GFP, three SCDs in *C. elegans*, were upregulated by ZnSO_4_ treatment while downregulated by TPEN, the levels of oleic acid C18:1(n-9) and the conversion activity of SCD (C18:1(n-9)/C18:0) were increased under TPEN treatment, but were reduced in *cdf-1(n2527)* and *sur-7(tm6523)* mutants by ZnSO_4_ treatment. Thus, these results demonstrate that a negative regulation of SCD expression and conversion activity by zinc, further confirms our hypothesis that zinc acts bypass SREBP to directly determine the conversion activity of SCD in lipid biosynthesis and accumulation. Therefore, we speculate that targeting of SCD may provide potential possibility for the treatment of zinc-related lipid metabolic diseases.

## Acknowledgements

This work was supported by the National Natural Science Foundation of China (31671230, 91857113, 81700520, 31860323, 31801001, U1702288, U1702287), the Ministry of Science and Technology of the People’s Republic of China (2018YFA0800700), the Yunnan Applied Basic Research Projects (2017FA007, 2018FB117, 2019FB046, 2019FB048), and Yunnan Oversea High-level Talents Program (2015HA039 and 2015HA040 to BL).

## Author Contributions

Y.H. and J.J.Z. conceived and designed the experiments, and wrote the paper. Y.L.W. revised the manuscript. YH. and J.J.Z. performed most of the experiments and data analysis, Y.L.W., L.F., X.Y.L and YW performed some experiments. J.S., X.J.W. and X.Y.W provided the guidance for some experiments and contributed reagents/materials/analysis tools. All authors reviewed the manuscript.

## Supplementary information

**Figure S1.**
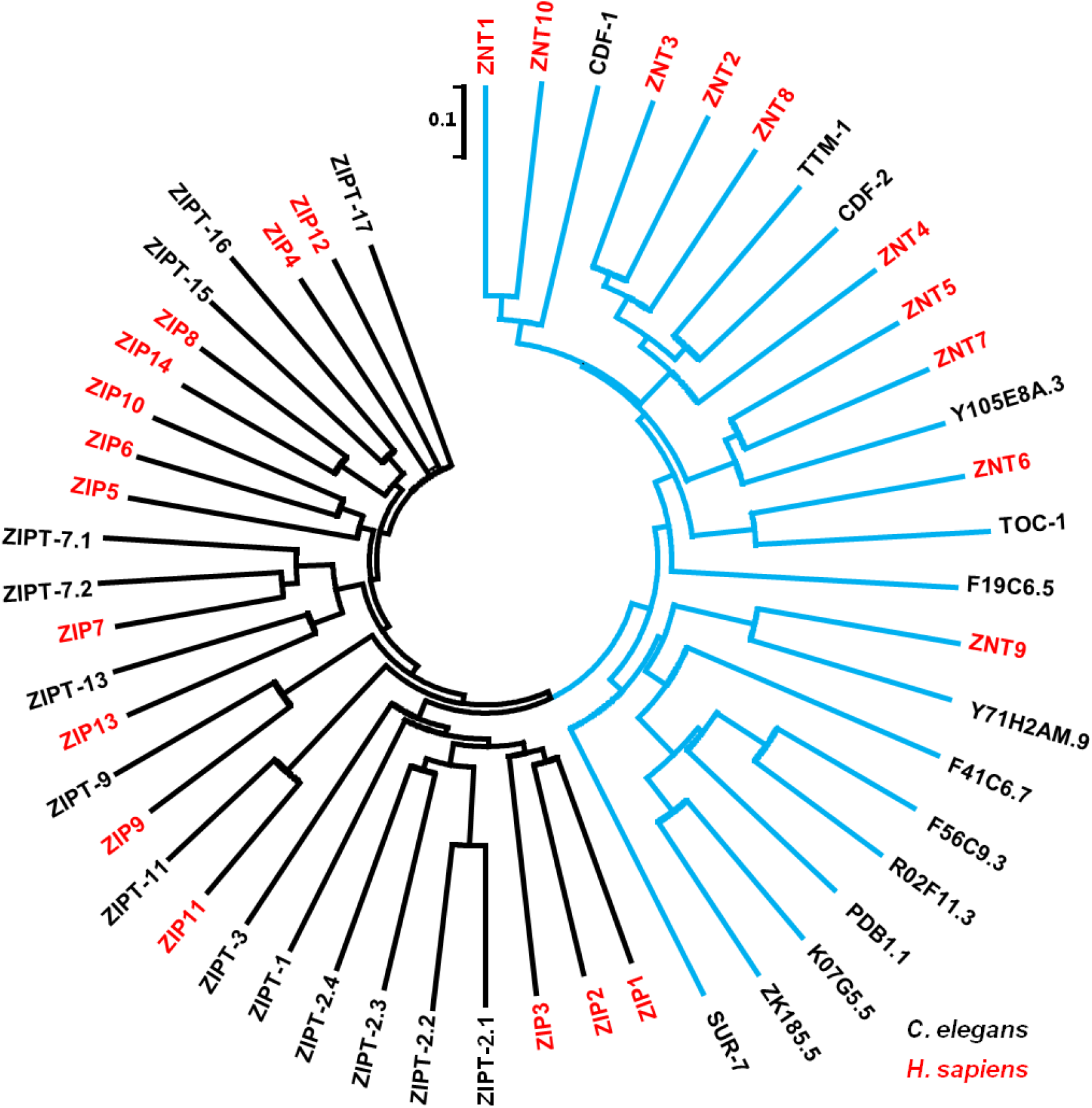
Zinc transport relevant proteins cladogram. A dendrogram shows zinc transport relevant proteins from *Caenorhabditis elegans* (28 members, black typeface) and *Homo sapiens* (red typeface), including CDFs family (blue line) and ZIPs family (black line).

**Figure S2.**
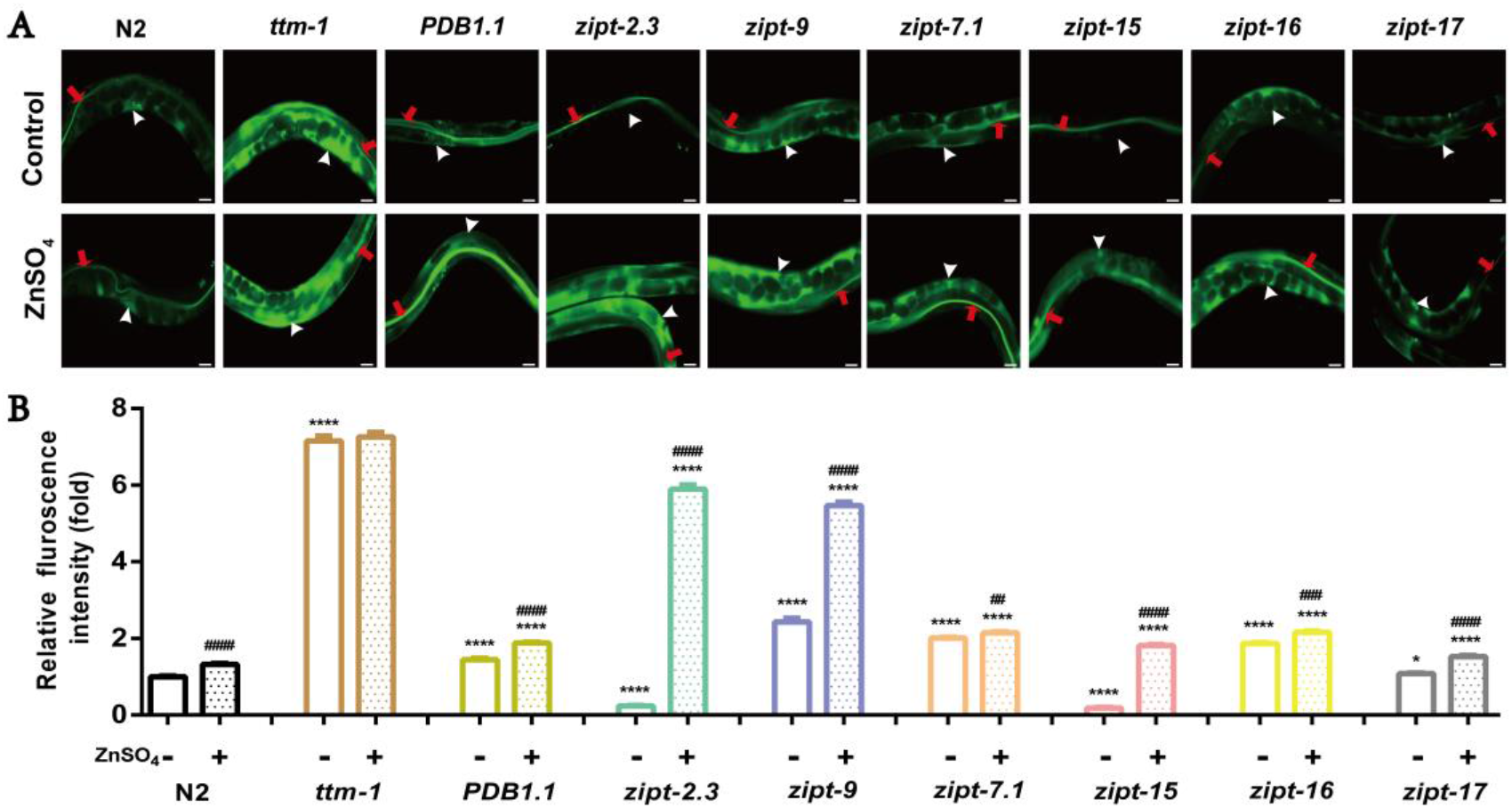
Other zinc related transporter mutants with zinc content. (A) Fluorescence microscopy of live worms of N2 (WT), *ttm-1(tm6669), PDB1.1(ok3114), zipt-2.3(ok2094), zipt-9(ok876), zipt-7.1(ok971), zipt-15(ok2160), zipt-16(ok875)* and *zipt-17(ok745)*, which stained with zinpyr-1 and captured using identical settings and exposure times. Scale bar represents 25 μm. The concentration of ZnSO_4_ is 50 μM. The red arrowheads indicate the lumen zinc accumulates of the intestine, the white arrows show pseudocoelomic zinc. (B) Quantitation of the fluorescence intensity about pseudocoelomic zinc.

**Figure S3.**
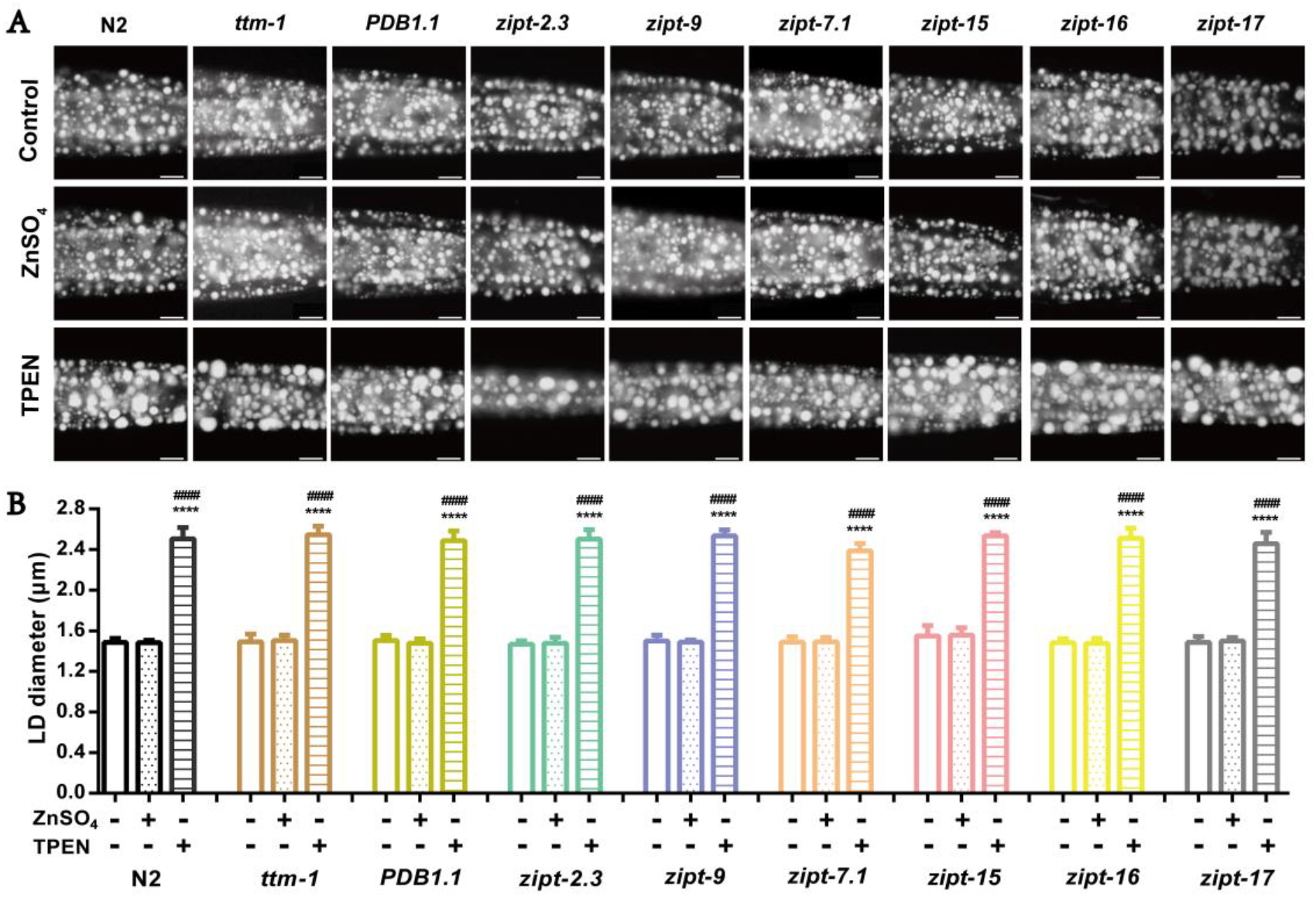
The fat content of other zinc related transporter mutants in specific zinc condition. (A) Nile red staining of fixed worms of N2(WT), *ttm-1(tm6669), PDB1.1(ok3114), zipt-2.3(ok2094), zipt-9(ok876), zipt-7.1(ok971), zipt-15(ok2160), zipt-16(ok875)* and *zipt-17(ok745)* under control, ZnSO_4_ and TPEN treatment. For representative animals, the anterior was on the left and the posterior was on the right, Bar, 10 μm. (B) Quantitation of lipid droplet size in the posterior region of intestines from five worms of each worm strain.

**Figure S4.**
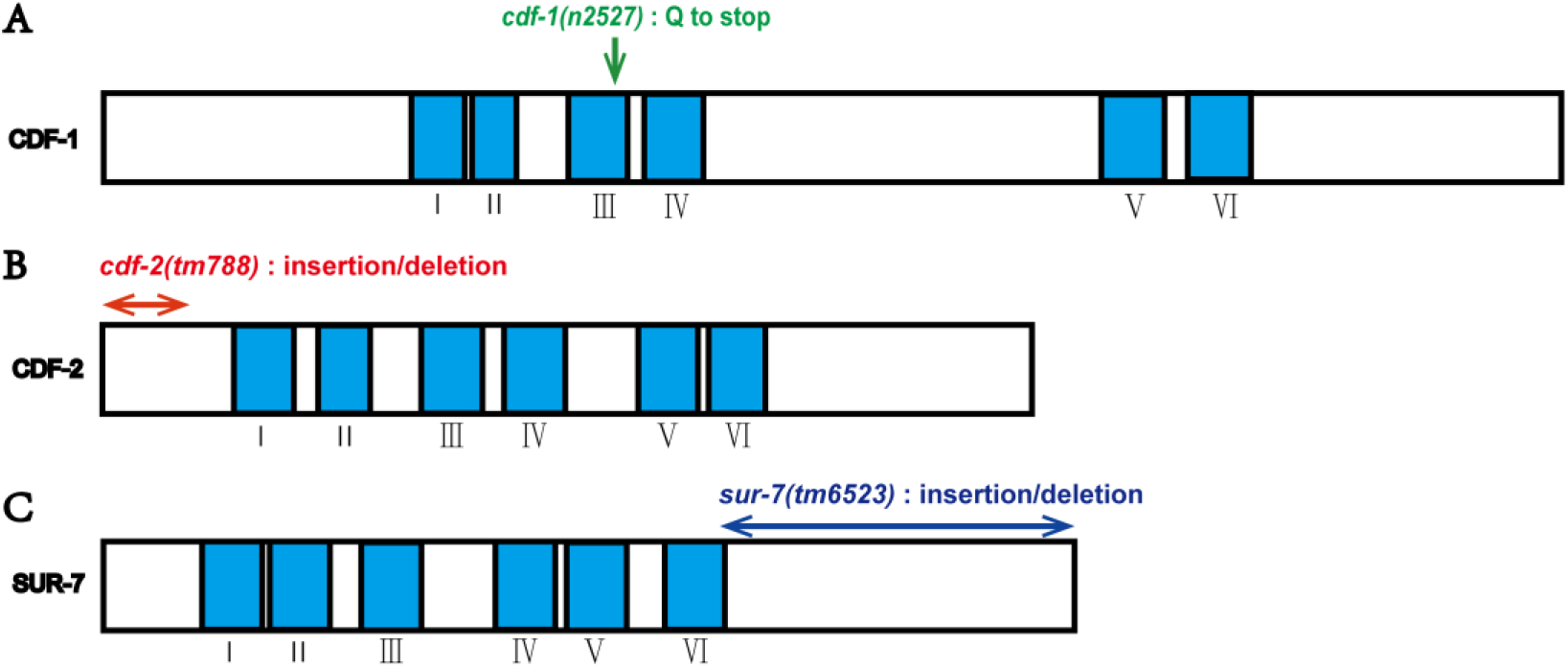
The structure of (A) CDF-1, (B) CDF-2 and (C) SUR-7 protein domains. The blue rectangles indicate the predicted six transmembrane motifs (labeled I–VI). (A) The green arrow indicates the stop codon replaced amino acids Q of *cdf-1(n2527)* mutant. (B) The double red sided arrow indicates the insertion/deletion region, which cause frame shift mutation of *cdf-2(tm788)*. (C) The double blue sided arrow indicates the insertion/deletion region, which cause early termination of translation of *sur-7(tm6523)* mutant.

**Figure S5.**
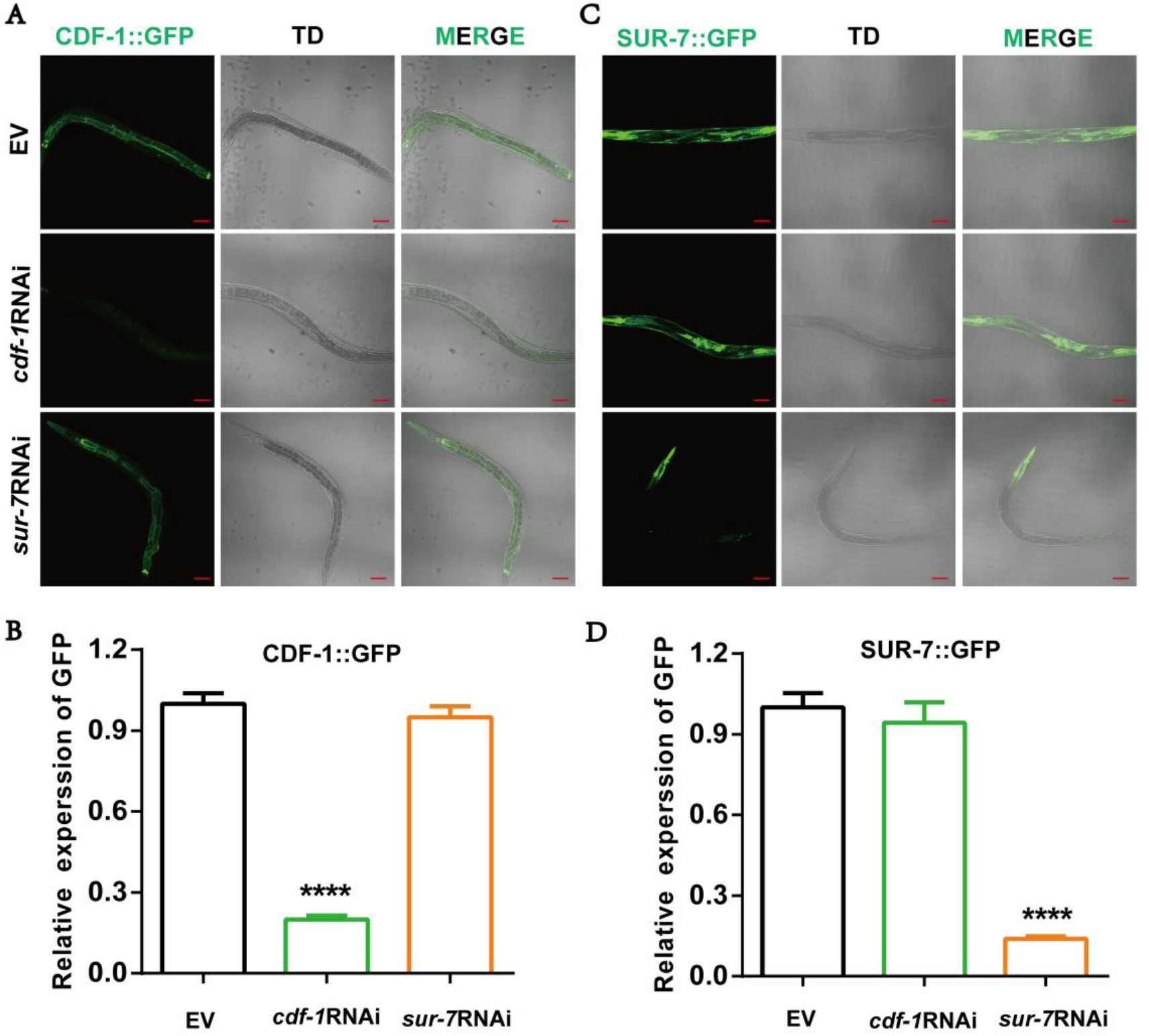
Validation of RNAi efficiency and specification. (A) Fluorescence intensity of CDF-1::GFP under empty vector EV, *cdf-1*RNAi and *sur-7RNAi*. Bar, 100 μm. (B) Quantization of fluorescence intensity of CDF-1::GFP. (C) Fluorescence intensity of SUR-7::GFP under empty vector EV, *cdf-1*RNAi and *sur-7RNAi*. Bar, 100 μm. (D) Quantization of fluorescence intensity of SUR-7::GFP.

**Table S1.**
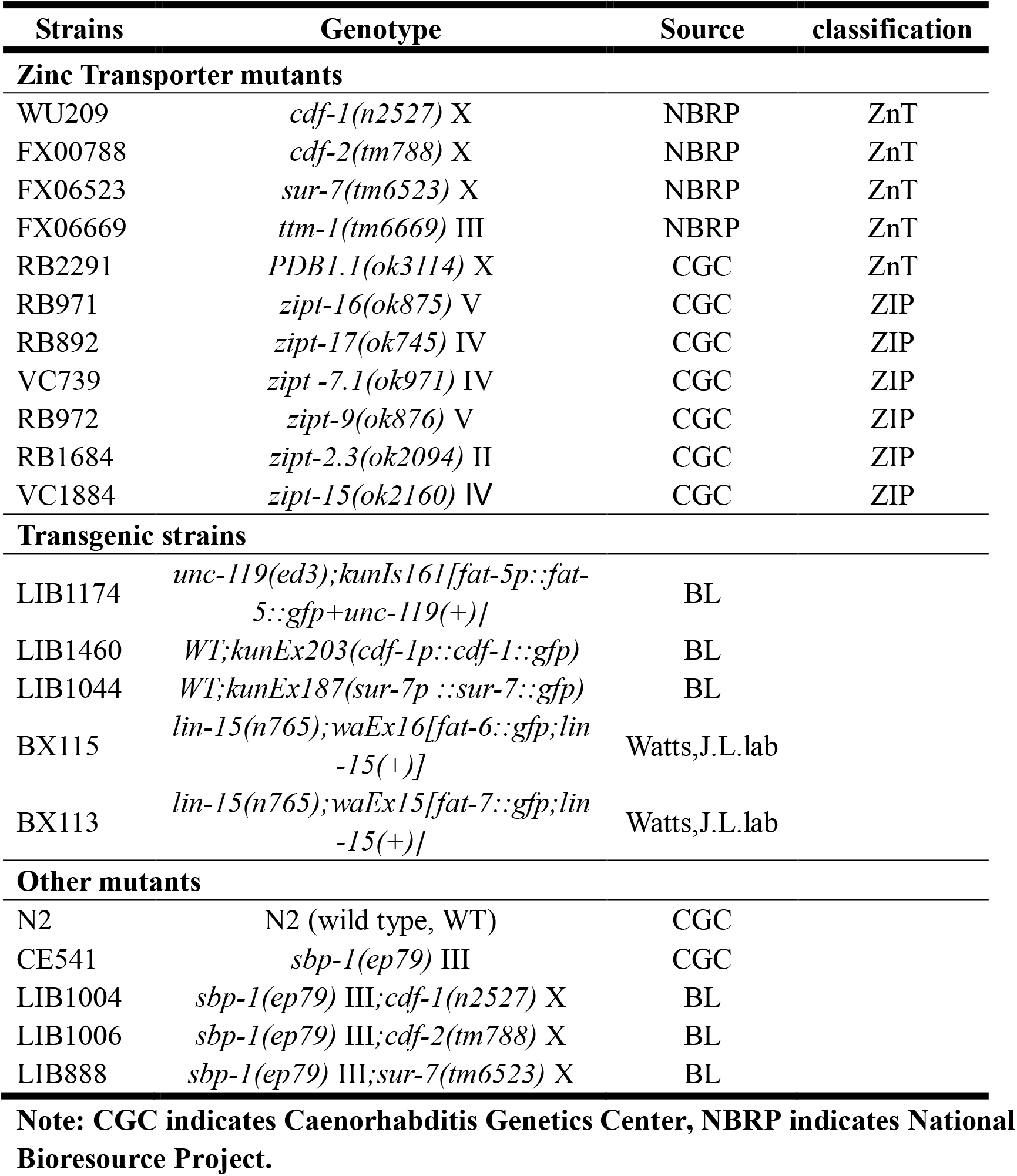
Information of worm strains used in this study

| Strains | Genotype | Source | classification |
| --- | --- | --- | --- |
| **Zinc Transporter mutants** |  |  |  |
| WU209 | *cdf-1(n2527)* X | NBRP | ZnT |
| FX00788 | *cdf-2(tm788)* X | NBRP | ZnT |
| FX06523 | *sur-7(tm6523)* X | NBRP | ZnT |
| FX06669 | *ttm-1(tm6669)* III | NBRP | ZnT |
| RB2291 | *PDB1.1(ok3114)* X | CGC | ZnT |
| RB971 | *zipt-16(ok875)* V | CGC | ZIP |
| RB892 | *zipt-17(ok745)* IV | CGC | ZIP |
| VC739 | *zipt-7.1(ok971)* IV | CGC | ZIP |
| RB972 | *zipt-9(ok876)* V | CGC | ZIP |
| RB1684 | *zipt-2.3(ok2094)* II | CGC | ZIP |
| VC1884 | *zipt-15(ok2160)* IV | CGC | ZIP |
| **Transgenic strains** |  |  |  |
| LIB1174 | *unc-119(ed3);kunIs161[*fat-5p::fat-5::gfp+unc-119(+)*]* | BL |  |
| LIB1460 | *WT;kunEx203(*cdf-1p::cdf-1::gfp*)* | BL |  |
| LIB1044 | *WT;kunEx187(*sur-7p::sur-7::gfp*)* | BL |  |
| BX115 | *lin-15(n765);waEx16[*fat-6::gfp;lin-15(+)*]* | Watts,J.L.lab |  |
| BX113 | *lin-15(n765);waEx15[*fat-7::gfp;lin-15(+)*]* | Watts,J.L.lab |  |
| **Other mutants** |  |  |  |
| N2 | N2 (wild type, WT) | CGC |  |
| CE541 | *sbp-1(ep79)* III | CGC |  |
| LIB1004 | *sbp-1(ep79)* III;*cdf-1(n2527)* X | BL |  |
| LIB1006 | *sbp-1(ep79)* III;*cdf-2(tm788)* X | BL |  |
| LIB888 | *sbp-1(ep79)* III;*sur-7(tm6523)* X | BL |  |

**Note: CGC indicates Caenorhabditis Genetics Center, NBRP indicates National Bioresource Project.**

**Table S2.**
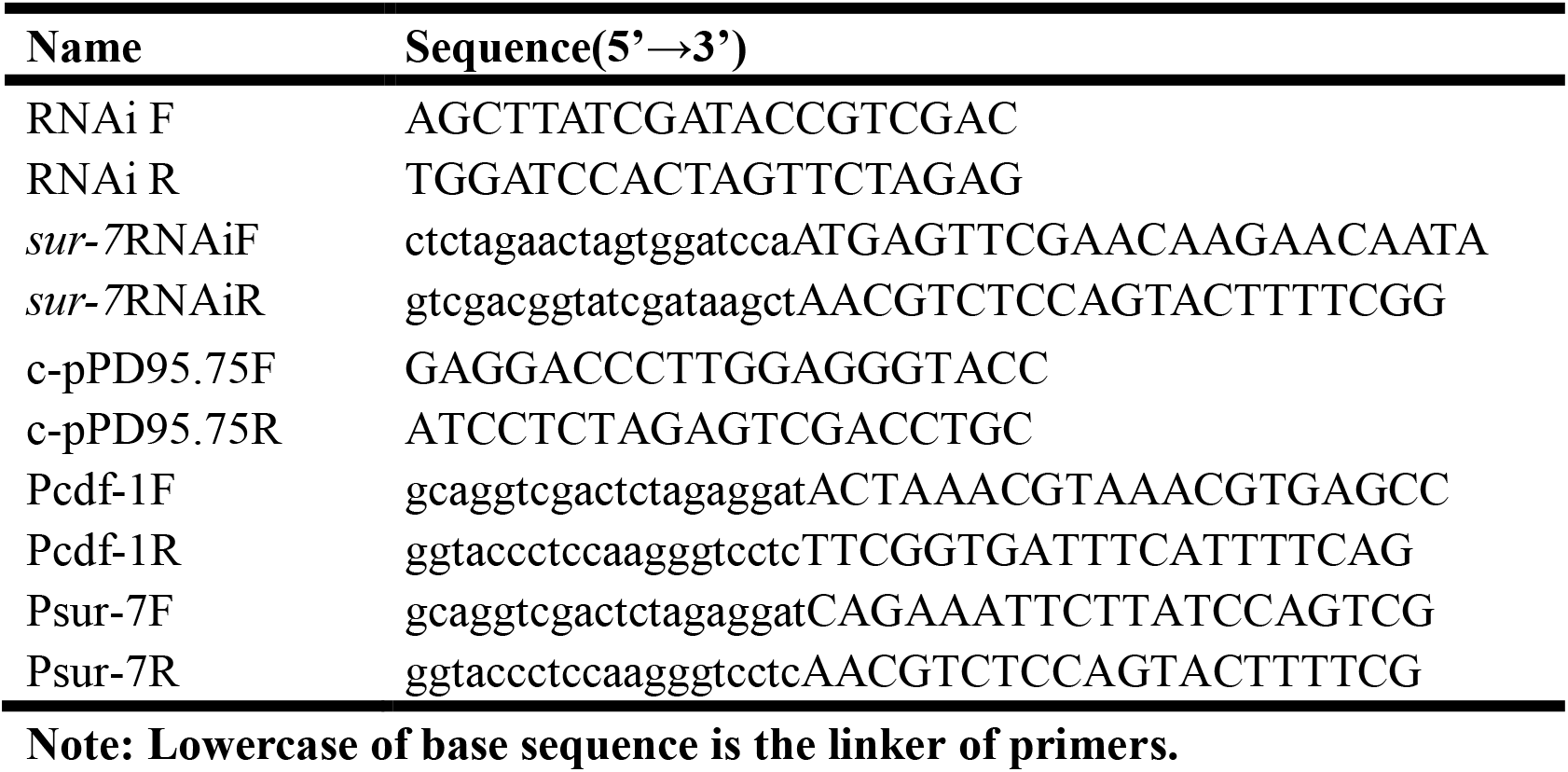
Primers

